# Testosterone induces a female ornament followed by enhanced territoriality in a tropical songbird

**DOI:** 10.1101/2020.04.27.065011

**Authors:** Jordan Boersma, Erik D. Enbody, John Anthony Jones, Doka Nason, Elisa Lopez-Contreras, Jordan Karubian, Hubert Schwabl

## Abstract

We know little of the proximate mechanisms underlying expression of signaling traits in female vertebrates. Across males the expression of sexual and competitive traits, including ornamentation and aggressive behavior, is often mediated by testosterone. In the White-shouldered Fairywren (*Malurus alboscapulatus*) of New Guinea, females of different subspecies differ in presence or absence of white shoulder patches and melanic plumage, while males are uniformly ornamented. Previous work has shown that ornamented females circulate more testosterone and exhibit more territorial aggression than do unornamented females. We investigated the degree to which testosterone regulates expression of ornamental plumage and territorial behavior by implanting free-living unornamented females with testosterone. Every testosterone-treated female produced a male-like cloacal protuberance, and 15 of 20 replaced plucked brown feathers with white shoulder patch feathers, but did not produce melanic plumage characteristic of ornamented females. Testosterone treatment did not elevate territorial behavior prior to production of the plumage ornament and exhaustion of the implant. However, females with experimentally induced ornamentation, but exhausted implants, increased the vocal components of territory defense relative to fully unornamented control and also to testosterone-implanted females. Our results suggest that testosterone induces partial acquisition of the ornamental plumage phenotype, and that ornament expression, rather than testosterone alone, results in elevated territorial behavior.

**Lay Summary:** Testosterone regulates expression of a suite of competitive traits in male organisms and could have similar function in females. Empirical tests are needed to determine the extent to which testosterone promotes production of ornamentation and competitive behaviors in female animals. We supplemented testosterone in unornamented females of a species where naturally occurring ornamented females circulate higher testosterone and are more territorially aggressive. Implanted females produced partial ornamentation, which was followed by increased territoriality that was apparently unrelated to testosterone circulation itself.

## INTRODUCTION

The expression of elaborate secondary sexual traits in females, such as ornaments, was long thought to be a byproduct of sexual selection on males (Darwin 1871). However, evolutionary transitions in elaborate coloration, or ornamentation, occur more frequently in females than in males in many taxa, suggesting that selection can act on female ornament evolution independently of selection on males (Irwin 1994; Omland 1997; Burns 1998; Johnson et al. 2013; Price and Eaton 2014). Moreover, recent empirical studies provide support for adaptive functions of ornamentation in females, sometimes in the context of sexual selection (eg. Fitzpatrick and Servedio 2017) or in the context of competing for non-mating resources, via social selection (West-Eberhard 1979; West-Eberhard 1983; reviewed in Tobias et al. 2012). One useful route to evaluating the function of a putative signal is to understand the proximate mechanisms regulating its expression (Hau 2007; Rosvall et al. 2016), yet very little is known about mechanisms of ornament production in females.

Differential secretion of sex steroids is a common mechanism underlying sex-specific trait expression from a shared autosomal genome. For instance, in males of some avian taxa, increased circulation of androgens induces molt into colorful male plumage, while in other taxa enhanced estrogen circulation in females induces molt into cryptic female plumage (reviewed in Kimball and Ligon 1999). Sex steroids like testosterone mediate development and expression of a suite of morphological and behavioral traits (reviewed in Hau 2007), thus causing some evolutionary endocrinologists to label testosterone as a phenotypic integrator (reviewed in Lipshutz et al. 2019). There is some empirical support for testosterone acting as a phenotypic integrator in males (Ketterson and Nolan Jr 1999; Wingfield et al. 2001; Hau 2007a; Hau and Wingfield 2011), though it can be difficult to determine how traits are linked mechanistically due to the dynamic feedback between hormones and the traits they potentially regulate (Rubenstein and Hauber 2008; Safran et al. 2008; Vitousek et al. 2014).

The role of testosterone in female ornamentation is currently equivocal. However, testosterone is correlated with female ornamentation in several bird species (Muck and Goymann 2011; Moreno et al. 2014; Cantarero et al. 2017), and exogenous testosterone has been found to stimulate production of ornament production in females of several avian and non-avian species (Lank et al. 1999; Eens et al. 2000; Lahaye et al. 2012; Cox et al. 2015; Lindsay et al. 2016). Testosterone supplementation can also stimulate competitive behaviors in females including singing (Kriner and Schwabl 1991; De Ridder et al. 2000) and territorial aggression (Zysling et al. 2006; Rosvall 2013; Cantarero et al. 2015). However, most testosterone manipulation experiments have been conducted in species lacking discrete variation in female phenotypes, so these studies, while useful for determining mechanisms of sexual dimorphism and capacity for phenotypic plasticity in these systems, do not inform our understanding of the proximate basis of naturally varying female phenotypes.

In the current study, we supplemented testosterone to female White-shouldered Fairywrens (*Malurus alboscapulatus*, Maluridae), a passerine bird endemic to New Guinea. In this species, females show pronounced variation in plumage ornamentation by subspecies while males are similarly ornamented across subspecies. Previously, we found that females belonging to the subspecies (*M. a. moretoni*) with contrasting black-and-white female ornamental plumage have higher mean plasma testosterone levels and also show greater territorial defense behavior than females belonging to the subspecies lacking this ornamentation (*M. a. lorentzi*; Enbody et al. 2018), suggesting that testosterone might play a role in female trait variation. Males from a closely related species, the Red-backed Fairywren (*Malurus melanocephalus*), provide a useful comparison to our system, as they vary from cryptic female-typical brown plumage to red-and-black ornamental plumage. Ornamented males circulate higher testosterone and are more aggressively territorial than unornamented males (Barron et al. 2015), and exogenous testosterone stimulates the acquisition of ornamental plumage in males (Lindsay et al. 2011). Females in this species are naturally unornamented, but nonetheless developed a partial, male-like phenotype when implanted with testosterone (Lindsay et al. 2016). Collectively, the results in Red-backed Fairywrens inform the prediction that the ornamented female White-shouldered Fairywren phenotype is produced via enhanced testosterone circulation.

Here, we experimentally assess whether testosterone regulates expression of ornamentation and the associated behavioral phenotype in a natural population of a female vertebrate. Second, we investigate the links between testosterone, ornamentation, and territorial behavior. Third, we assess whether expression of ornamentation generates a social cost in the form of increased reception of territorial aggression from conspecifics. Specifically, we address three hypothesized pathways to expression of the integrated ornamented female phenotype: 1) exogenous testosterone supplementation in unornamented females of *M. a. lorentzi* induces molt into the ornamented plumage phenotype observed in *M. a. moretoni*; 2) testosterone also promotes territory defense behaviors, independently of plumage ornamentation; 3) expression of ornamental plumage promotes territory defense independent of elevated testosterone.

## METHODS

### Study system and general field methods

We studied White-shouldered Fairywrens (Figure 1A., *Malurus alboscapulatus lorentzi*) in Obo village, Western Province, Papua New Guinea (Figure 1B., 141°19′ E, 7°35′ S, 10–20 m a.s.l.).

**Figure 1:**
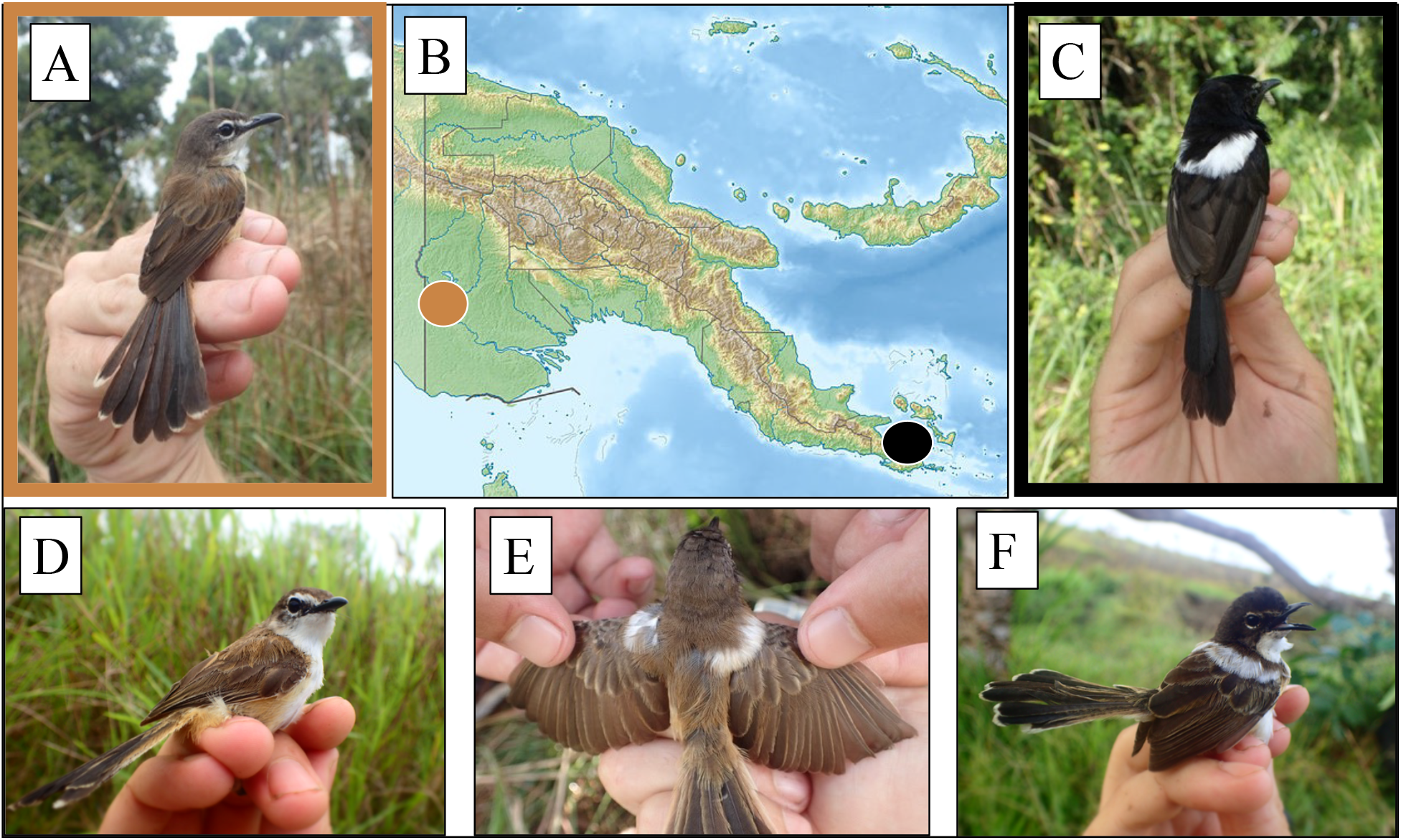
**A.** An unmanipulated female from the unornamented White-shouldered Fairywren subspecies (*Malurus alboscapulatus lorentzi*). **B.** Map of Papua New Guinea depicting the location of our two field sites (brown dot: field site of *M. a. lorentzi*, where our study was conducted; black dot: field site of *M. a. moretoni*). **C.** An unmanipulated ornamented female from the *M. a. moretoni* subspecies. **D-F.** Testosterone-implanted females of *M. a. lorentzi* displaying a gradient of ornamentation: **D.** no ornamentation (N = 5), **E.** partial shoulder patches (N = 11) **F.** prominent shoulder patches with darker brown body plumage (N = 4).

Females of this subspecies lack the black plumage with white shoulder patches that is expressed in males of this species as well as females of other subspecies (e.g., *M. a. moretoni*; Fig. 1C; Enbody et al. 2018). Females were captured predominantly by flushing into mist nets, and in rarer cases, using duet playback. Individuals who were previously unbanded were given a unique Australian Bird and Bat Banding Scheme metal band in addition to a unique combination of three plastic color bands. White-shouldered Fairywrens nest year-round (Enbody et al. 2019), so in order to mitigate the confounding effects of breeding stage on behavior we excluded females who were actively nesting from the experiment.

### Experimental design

#### Testosterone implants

Each implant consisted of 14.63 mg of beeswax (Beesworks®), 4.87 mg peanut oil (ACROS Organics™) and 0.5 mg of crystalline testosterone (Sigma T1500), initially dissolved in 2.5 μl 100% ethanol (Fisher Bioreagants™). Control implants were identical in composition to the testosterone implants except that they lacked testosterone. Testosterone and control implants were both roughly 2×3.2mm (diameter x length) and weighed between 19.8 and 20.7mg. We scaled implants to produce physiologically maximal levels for White-shouldered Fairywren females (Khalil et al., in review; H. Schwabl and W. Goymann, unpublished data). Each implant was inserted subcutaneously in the abdominal region after plucking feathers from the area and sterilizing the incision site with rubbing alcohol, then the incision was sealed with VetBondTM (3M). We implanted 4 females with testosterone implants and 4 with control implants during a pilot study in September 2016 and 16 females with testosterone and 8 with control implants in March 2017. All assessments of behavior were conducted in 2017.

#### Plumage, molt, and cloacal protuberance assessments

We plucked approximately 10 feathers at initial capture (when implants were set) across 4 body regions: crown, shoulder patch, rump, chest, and 2 tail feathers to induce feather replacement. Ornamental female plumage is characterized by black feathers on the crown, rump, chest, and tail, and white shoulder patch feathers (Enbody et al. 2017). During the 2016 pilot study we captured implanted females approximately 1, 2, 3, and 4 weeks post-implanting to assess the effect of testosterone on feather replacement, ornamentation, and morphology. In 2017 we captured a subset of females around 10 days after implanting (n = 11 testosterone females; 6 control females) and again 28+ days later (n = 11 testosterone females; 5 control females) to align our morphological measurements to the behavioral experiments described below. We assessed testosterone-induced cloacal protuberance size by measuring the length, width, height with digital calipers. Cloacal protuberance (CP) volume was estimated as volume = π (D/2)W/2))L (Tuttle et al. 1996). At each capture, molt and plumage were assessed across the following body regions: head, back, chest, belly, wing, and tail. Plumage coloration was qualitatively assessed by noting presence or absence of black or white feathers, and molt was quantified by determining the approximate proportion of actively molting feathers in each region on a scale of 0-3 (0 = no molt, 1 = 0-32%, 2 = 33-66%, 3 = 67-100% feathers molting).

#### Simulated territorial intrusions

We used an established simulated territorial intrusion (STI) protocol for our study system (detailed methods in Enbody et al. 2018). Briefly, we placed cardstock mounts painted to resemble pairs from *M. a. lorentzi* on a focal pair’s territory and lured in the pair using a duet recorded from our study population. We randomly selected one male and one female mount (n = 4 mounts for each sex) and a duet (n = 10) for each trial; in the rare case that duets were from the focal pair or their territory neighbors we re-selected a different duet. Each duet was played through a small speaker (UE Roll 2; California, USA) and consisted of a single duet separated by 10 seconds of silence before repeating. Mounts were placed immediately next to each other above the speaker on a 1.5m tall stick containing three small branches: one containing the mount pair, and the others for responding pairs to perch on. Though our main goal was to quantify female response, we used mount pairs and duets for the following two reasons: because we have never observed White-shouldered Fairywren females invading territories on their own, and because using only a female mount and song can potentially lead to multiple males approaching to perform sexual displays to the mount (Enbody and Boersma, personal obs.).

We started each trial with 1 min of acclimation time followed by 5 min of duet playback, or until the focal female approached within 1m of the mount, whichever occurred first. In the former case, we played 3 more duets before ceasing playback, and in both cases, we continued to record behavior until 5 minutes after the playback was stopped. The goal of this approach was to standardize the number of duets each responding pair was exposed to while in close proximity to the speaker. If the pair or the focal female failed to approach within 10m of the mount after 5 minutes the trial was terminated, we searched the territory for the pair and started a new trial close to their current location. We recorded data for 2 female responses where the male did not approach within 10m. The behavior of territory-holding males and females (pairs) was quantified separately by one observer 20m away from the mounts. For each we quantified the rate of the following behaviors: duets, solo songs, flybys (where an individual flew within 1m of the mount), and the proportion of time within 5m of the mount. Additionally, we recorded latency to first vocalization (duet or solo song) and to approach within 5m of mounts.

The design of our STI experiment in 2017 was informed by pilot work in 2016 where we found that exogenous testosterone induced partial ornament expression in females. We repeated STI trials a maximum of 3 times for each pair included in the female implant experiment: (1) prior to implanting; (2) 5-10 days after implanting with testosterone or control; (3) 28+ day after implanting (Fig. 2). At time point 2, females implanted with testosterone were expected to have levels elevated well beyond their baseline (time point 1), but have yet to develop plumage ornamentation, and at time point 3 testosterone from the implant should have been exhausted (H. Schwabl and W. Goymann, unpublished data) and ornamental feathers have fully emerged from their sheaths and readily visible. This approach allowed for us to test both how testosterone and ornament presence influenced territorial aggression independently.

**Figure 2:**
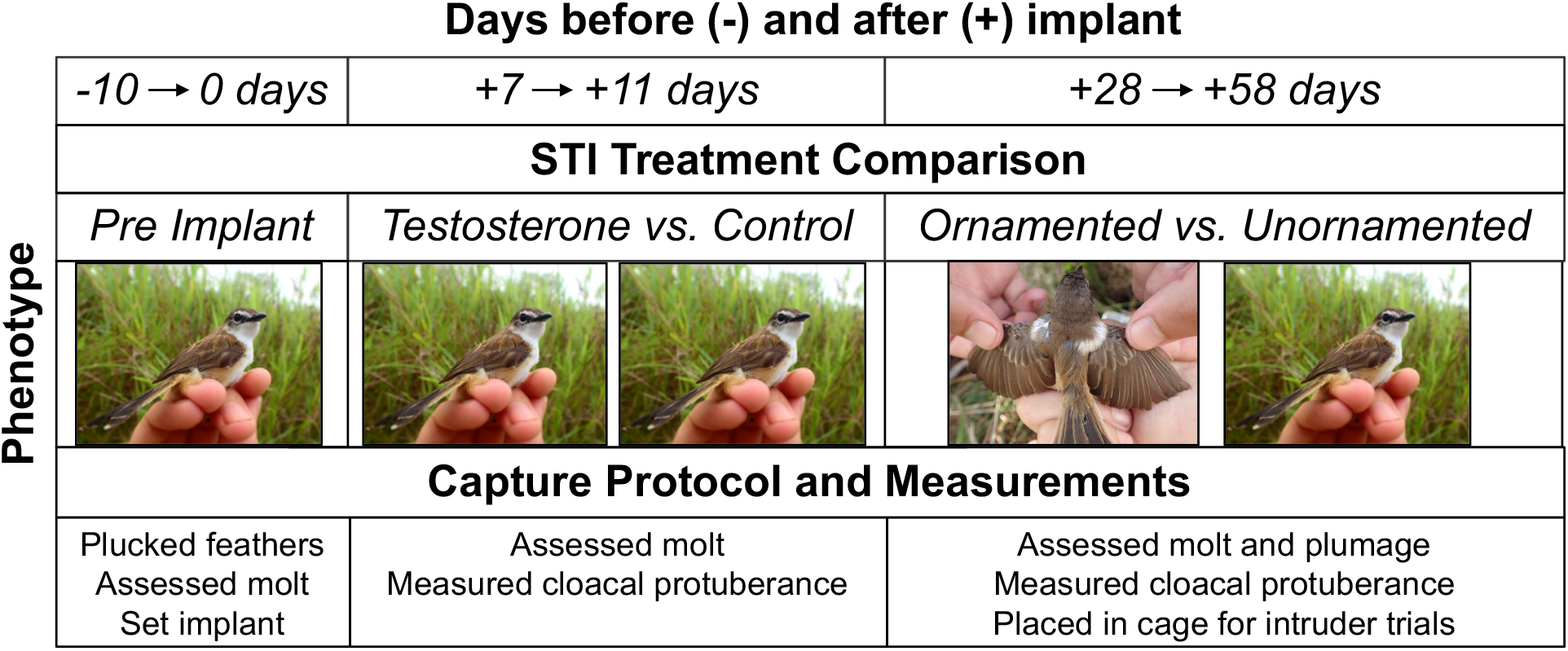
Experimental timeline for STI and morphological sampling. Photographs for phenotype simply reflect general physical appearance for each treatment type rather than unique photographs from each treatment type during that sampling period. Females only differed in plumage during the final sampling period (28-58 days post-implant).

#### Caged female intrusions

We designed a second behavioral assay to assess how presence of ornamentation influences the territorial defense behaviors of conspecifics. Following the general methods of Karubian et al. (2008), we used live females belonging to either the experimentally induced ornamented phenotype or natural unornamented phenotype as stimuli and assessed the response of free-living pairs. We built a 63.5 cm^3^ cage using a wood frame and wire mesh with two small bamboo sticks for perching within the cage (Figure S3). At the bottom of the cage was a cloth bird bag that opened into the cage so we could release the female and the bag was cinched closed during the trial to ensure that the caged individual could not escape. The cage was placed toward the center of a known White-shouldered Fairywren territory using 2 large sticks that held the cage ~1.5m above the ground (Figure S3). We chose a microhabitat that would allow for the cage to sit near the top of the grass in order to mimic where this species is often found in its habitat and also to allow for the bird to be easily visible to conspecifics. In order to minimize the time females spent in cages we did not include any acclimation time before proceeding with behavioral trials. These trials were conducted about 43 – 56 days after setting testosterone or control implants.

We used an amended version of our STI protocol for cage trials. As per the STIs, we played duet songs to lure the territory-holding pair to the area, but once they were within 5m we immediately ceased playback and then recorded their response for 10 min. During preliminary trials we determined that pairs apparently noticed the caged individual once they were within 5m of the cage and then directed their territory defense at the caged individual. Because we were exclusively interested in assessing response to the caged female’s state of ornamentation, we stopped the vocal stimulus of playback once pairs were close enough to perceive their intruder. If pairs failed to approach the cage within 5m after 5 min of playback the trial was terminated, and we moved to a neighboring territory. We quantified the same behavior rates as for the STIs but analyzed time within 0.5m of the cage instead of 5m. We did not observe any males displaying to the caged females, and both members of the pair usually responded to the intruding female with changes in behavior once within 5m. A camera was used to quantify the activity level of the caged stimulus female to determine if her behavior might affect pair response in addition to her plumage phenotype. We quantified caged female activity on a four-point scale: a score of 0 indicated no movement at all, 1 corresponded to movement while the territorial pair was absent, still when they were present, 2 denoted movement during most of the trial, and 3 represented constant movement for the duration of the trial.

### Statistical analysis

#### Cloacal protuberance volume

We analyzed cloacal protuberance (CP) volume from repeated measures taken from females belonging to the testosterone-treatment group (untreated and control females did not produce a CP). The purpose of these analyses was to assess whether there was any evidence for differential testosterone levels among females producing ornaments versus those that did not. We used t-tests in R version 3.5.1 (R Core Team 2018) to compare cloacal protuberance volume between females who produced some ornamentation to those that failed to produce any ornamentation.

#### Response to simulated territorial intrusions

We scaled and centered all STI response variables, then used principal components analysis (PCA) with an oblique promax rotation to quantify male and female response individually. Sexes were run in separate PCAs due to the possibility of female treatment affecting response variably between sexes. We used the R package *psych* (v1.8.12, Revelle 2018) for both PCAs, and PCA scores were analyzed using linear-mixed models in the R package *lme4* (v1.1-21, Bates et al. 2015). Scree plots were used to determine which PCs to analyze (eigenvalues > 1.0, Fig. S2). We assessed normality of PC scores with a Shapiro-Wilk test prior to building models. Following detection of a significant effect of treatment on PC scores, we used post-hoc Tukey comparisons to determine which groups differed. We included individual ID (for both members of pair), mount stimulus (1-4 for both sexes), and duet stimulus (1-10) as random effects in the model. Pairs who were later determined to have an active nest during the trial were excluded from analyses (n = 2), as were females who lost their implant (n = 1) or had an equivocal state of ornamentation (e.g. only a few ornamented feathers; n = 2). In total we analyzed trials from 9 testosterone-implanted females and 4 controls during the pre-implant period, 10 testosterone-implanted and 5 controls 7-11 days post-implant, and 4 partially ornamented and 4 fully unornamented females 28+ days after implanting with testosterone.

### Response to caged female trials

Caged female trials were analyzed similarly to STIs. First, we scaled and centered response variables, then ran each sex individually in a PCA with a promax rotation. We used scree plots to select which components to analyze (Fig. S4). We then analyzed PCA scores using a linear-mixed model with individual ID and caged female activity scores as random effects.

## RESULTS

### Morphology

#### Plumage and molt

Fifteen of 20 testosterone-implanted *M. a. lorentzi* females produced white shoulder patch feathers, which are not naturally produced in this subspecies (Fig. 1). Some of the white scapular feathers were brown at the base, while others were white throughout. Among the 15 females with white scapular feathers 8 molted a mix of white and brown feathers in this region and thus did not produce a full shoulder patch. None of the implanted females produced a full complement of black feathers but 4 females across both study years replaced light brown feathers with darker brown plumage and a few individual black feathers (Fig. 1F). None of the control-implanted females molted in white scapular feathers or any darker brown or black plumage.

#### Cloacal protuberances

All testosterone-implanted females developed cloacal protuberances in both study years within 7-11 days of treatment (Table S1). These cloacal protuberances resembled those in males of the species in most cases, however 4 individuals produced only the tip of a cloacal protuberance, thus received a volume of 0 after measurement. Females captured after 4 weeks (range 28 – 58 days) had greatly diminished cloacal protuberance volumes, consistent with circulating testosterone having returned to baseline levels (Fig. S1B). We did not observe measurable cloacal protuberances in any females sampled 30+ days after implantation (N = 9 females), suggesting that testosterone from implants was, as expected from pilot work (H. Schwabl and W. Goymann, unpubl. data), fully exhausted at this point. Cloacal protuberance volumes of females developing ornaments did not differ from those of females that did not at 7 – 11 (t_2.91_ = −0.047, p = 0.97) and 28+ days post-implant (t_6.04_ = −1.18, p = 0.28).

### Simulated Territorial Intrusion Response

#### Female response

We analyzed 43 total trials for female response. The first 2 PCs (eigenvalues > 1) cumulatively explained 58% of the behavioral variation (29% for both PC 1 and PC 2; Table 2). We interpret high scores for PC 1 as indicative of the motivation to quickly approach the mounts and sustain that close proximity; high scores for PC 2 reflect quicker and greater singing (solo songs and duets) in response to the stimulus (Table 1).

**Table 1:** Variable loadings for female response to simulated territorial intrusions.

| **Female STI Response** | **PC 1: Close to Mount** | **PC 2: Vocalizations** |
| --- | --- | --- |
| Standard deviation | 1.47 | 1.44 |
| Proportion of Variance | 0.29 | 0.29 |
| Latency to sing or duet | 0.12 | -0.97 |
| Latency to 5m | -0.77 | -0.03 |
| Song & duet rate | 0.15 | 0.72 |
| Time within 5m | 0.93 | -0.06 |
| Flyby rate | 0.02 | 0.05 |

**Table 2:** Variable loadings for females and males for caged female intrusions.

|  | **Females** | **Males** |
| --- | --- | --- |
|  | PC 1 | PC 1 |
| Standard deviation | 2.93 | 3.54 |
| Proportion of Variance | 0.59 | 0.71 |
| Latency to sing or duet | -0.88 | -0.72 |
| Latency to 0.5m | -0.92 | -0.83 |
| Song & duet rate | 0.77 | 0.92 |
| Time within 0.5m | 0.78 | 0.90 |
| Flyby rate | 0.30 | 0.30 |

Testosterone treatment did not have a significant effect on PC 1 or PC 2 prior to molt of shoulder patches (+7 to +11 days after implanting; PC 1: F_2,21.68_ = 1.88, p = 0.18; PC 2: F_2,14.70_ = 0.016, p = 0.98). However, there was a significant effect of presence of ornamentation on PC 2 during the final sampling phase (days 28+ after implanting; F_2,10.43_ = 9.79, p = 0.004), when T from implants should have been exhausted. A Tukey comparison revealed that partially ornamented females scored significantly higher relative to both the pre-implant period (p < 0.001) and compared to date-matched testosterone-implanted females lacking any ornamentation (p =0.036). We did not detect an effect of mount or duet stimulus used on PC scores.

#### Male response

We analyzed 44 total trials for male response. Behavior variables loaded differently for males (Table S2) compared to females (Table 1), with PC 1 explaining 48% of the variation and indicating a quick and sustained close approach to the mount together with a rapid and persistent vocal response. The eigenvalue for PC 2 was substantially lower than the value for PC 1 (1.12 and 2.40, respectively) and the main variable that loaded on that component, flyby rate, was a behavior that was not typically exhibited by males during these trials, so we excluded PC 2 from analysis.

Males did not differ in their response according to their female’s initial hormone treatment (F_2,25.15_ = 1.91, p = 0.17) or later state of ornamentation (F_2,10.41_ = 1.92, p = 0.19). As per female analyses, we did not detect an effect of mount or duet stimulus on PC scores.

### Territorial Response of Resident Pair to Ornamented versus Unornamented female intruder

PC loadings for responses of resident males and females were similar, with PC 1 explaining 59% of the variation in response for females and 71% of the variation for males (Table 2). Male and female PC 1 scores were positively correlated (Pearson’s r_10_ = 0.83, p = 0.0009). We interpret higher PC 1 scores as greater overall aggression (faster and longer approach, faster to sing and greater song rate) towards the caged female intruder. We did not find any effect of the female intruder’s state of ornamentation (ornamented versus unornamented; Fig. 4) on the response of the resident territorial female (F_1,9.54_ = 0.62, p = 0.45) or male (F_1,6.99_ = 0.0034, p = 0.96). Further, caged female intruder behavior (activity score) did not influence male or female territorial response (F_1,2.26_ = 2.59, p = 0.14).

## DISCUSSION

We assessed whether elevated circulating testosterone stimulates ornamentation and the associated behavioral phenotype in female White-shouldered Fairywrens (*Malurus alboscapulatus*), a species with discrete female plumage phenotypes that differ in ornamentation and territorial defense behavior across subspecies (Enbody et al. 2018; Enbody et al. 2019). We show experimentally that exogenous testosterone caused unornamented females to produce some plumage ornamentation (predominantly shoulder patches; Fig. 1E-F) and find evidence that its presence then enhances territorial defense behavior (Fig. 3). Our results suggest that testosterone is partly responsible for ornament production in females and that the ornamentation itself, in turn, alters their territorial behavior. We found no evidence that acquisition of this putative signal comes with a social cost in the form of enhanced territorial aggression received by the signal bearer. Furthermore, the mates of experimental females did not appear to modulate their response to a simulated intruder according to the female’s initial testosterone treatment or resulting plumage phenotype.

**Figure 3:**
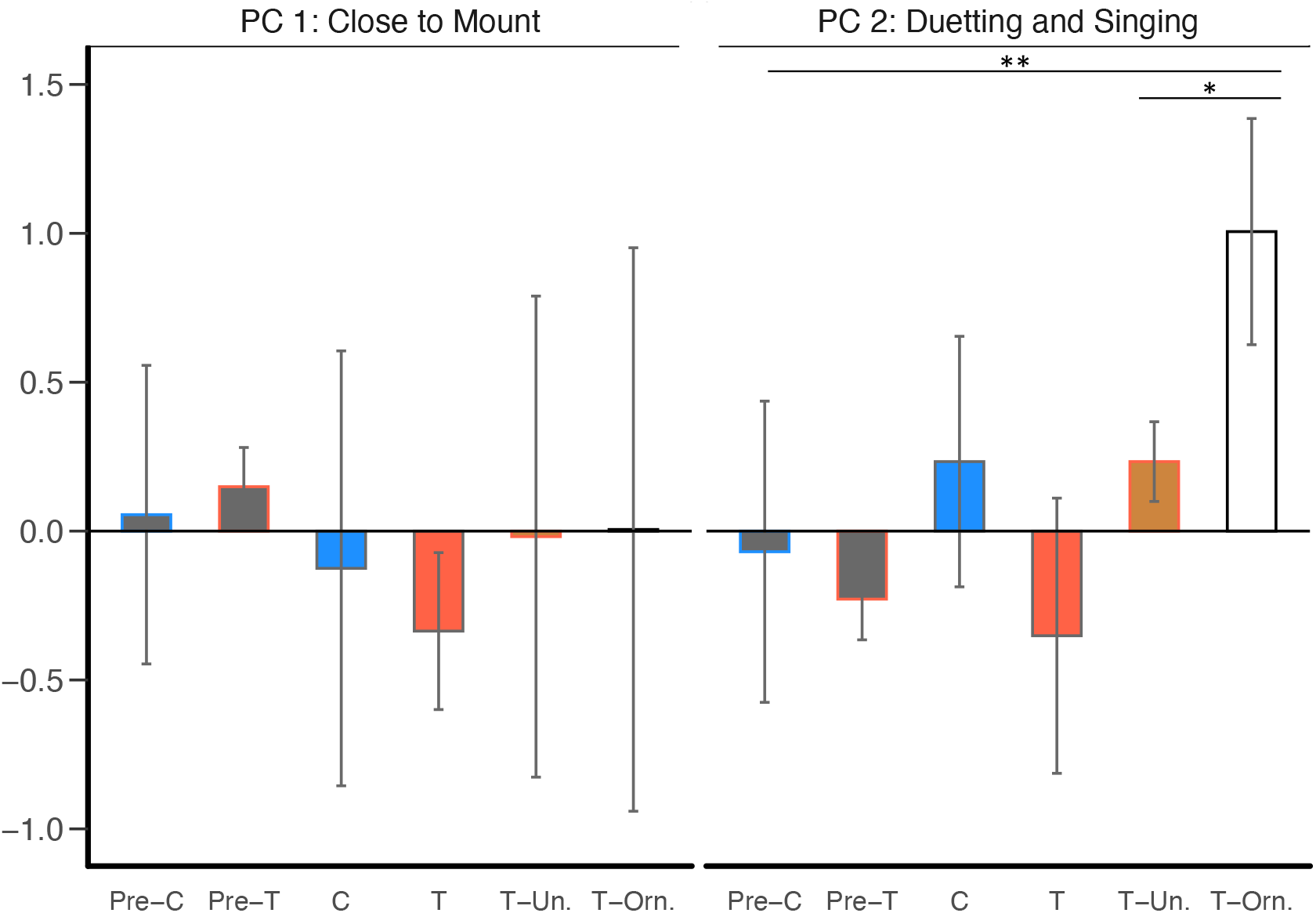
Female response to STIs by implant treatment, plumage phenotype, and progress of treatment. Females selected for testosterone or control treatment (pre-T; pre-C) did not initially differ in either PC. Testosterone treatment itself (7—11 days post-treatment) had no effect on either PC 1 or 2. Females who later (28—58 days post-treatment) had produced some ornamentation (T-Ornamented) had significantly greater PC 2 scores (vocal territoriality) compared to females within the testosterone-treated group who failed to produce ornamentation (T-Unornamented) and relative to the pre-treatment period. Bars denote means with plus/minus standard error lines overlaid; significant differences between treatment groups are indicated with asterisks (** p < 0.01, * p < 0.05).

This study is among the first to experimentally assess how testosterone mediates the production of elaborate female plumage within a species with naturally occurring alternate female phenotypes. The activational effects of testosterone on female plumage was mostly limited to acquisition of white components (shoulder patches) of ornamental plumage (Fig. 1E), and in rare cases darker brown feathers and a few black feathers in areas that are typically light brown (Fig. 1F). Our findings are consistent with results from a sister species lacking natural female ornamentation, the Red-backed Fairywren (*Malurus melanocephalus*), where testosterone-implanted females generated male-like, carotenoid-based red feathers and a darkened bill, but generally did not produce male-like melanic black feathers (Lindsay et al. 2016). In another member of the family Maluridae, Superb Fairywren (*Malurus cyaneus)*, testosterone-treated females underwent a male-like nuptial molt and feathers resembled males morphologically, however, females did not produce any of the elaborate coloration of the male ornamental plumage (Peters 2007). In two non-Malurid bird species exogenous testosterone led to enhanced bare part coloration in females (Eens et al. 2000; Lahaye et al. 2014), consistent with our plumage results. Finally, in reptiles, drab females typically respond to exogenous testosterone by producing male typical skin ornamentation matching, and in some cases, exceeding male skin coloration (reviewed in Cox et al. 2015). These studies and ours highlight that females often possess the mechanisms to respond to high circulating levels of testosterone but naturally do not express male-like ornamentation in part due to low circulating testosterone titers.

We found that elevated testosterone, before the production and expression of ornamentation, did not enhance territorial defense behavior (close approach and singing) during simulated territorial intrusions (STIs) relative to controls (Fig. 3). Exogenous testosterone increased aggressive behavior of females in many species studied to date (eg. Zysling et al. 2006; Sandell 2007; reviewed in Rosvall 2013). However, in some species testosterone-treatment did not enhance female aggressive behavior (DeVries et al. 2015) although it induced other androgen-regulated behavior such as singing in female European robins (*Erithacus rubecula*, Kriner and Schwabl 1991). Our results are consistent with an indirect relationship between circulating testosterone and aggression in White-shouldered Fairywrens, where testosterone promotes the production of a phenotype associated with aggression, but aggression is not elevated in the absence of that phenotype.

We found that females that had developed partial ornamentation in response to testosterone treatment showed greater singing and duetting during territory defense compared to females that did not produce ornamentation (Fig. 3). Previously a positive correlation between female ornamentation and aggression has been described in Gouldian Finches (*Erythura gouldiae*, Pryke 2007), in Pied Flycatchers (*Ficedula hypoleuca*, Morales et al. 2014), and in Lovely Fairywrens (*Malurus amabilis,* Leitão et al. 2019). Our results showing that females with testosterone-induced partial ornamentation exhibit a greater response during STIs (Fig. 3; PC2: territorial singing and duetting) are consistent with previous studies of effects of ornamentation on territorial behavior in male birds. Experimental enhancements of ornaments typically result in males achieving greater ranking within the social hierarchy via increased dominance in contests (reviewed in Vitousek et al. 2014). Our finding that females who acquired some ornamentation (shoulder patches) exhibited more vocal territoriality highlights the potential for shoulder patches acting as a status signal in intraspecific competition.

We believe that aggression is associated with plumage, but not testosterone for these reasons: 1) testosterone-implanted females lacking ornamentation did not differ behaviorally from controls when testosterone was elevated (Fig. 4), 2) testosterone-treated females displaying partial ornamentation differed from other testosterone-implanted females lacking ornaments, 3) size of androgen-dependent cloacal protuberances did not differ among partially ornamented and unornamented females receiving the same testosterone-treatment (Fig. S1B), indicating that testosterone levels were similar between females who did and did not produce ornamentation.

**Figure 4:**
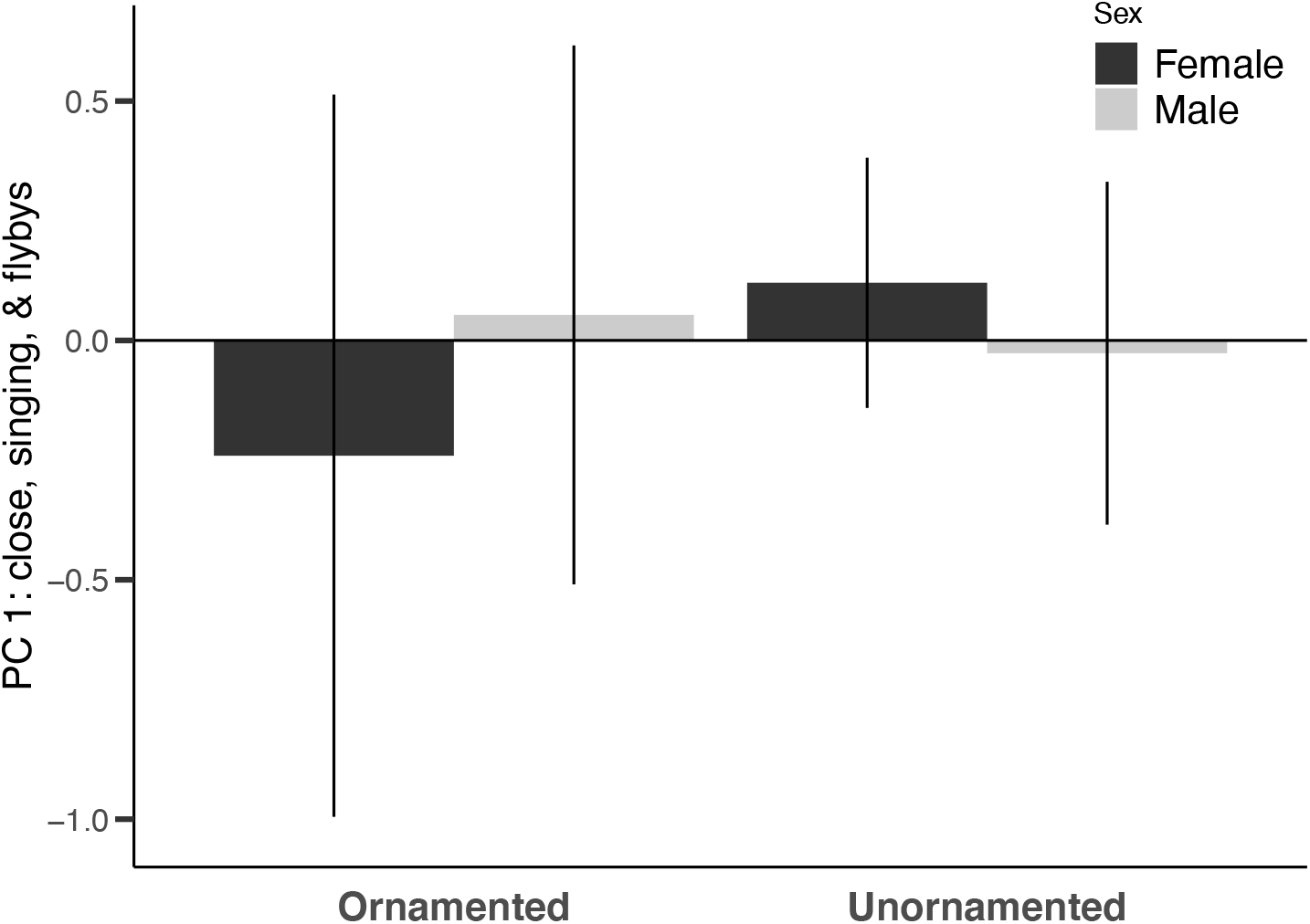
Response of resident females and males (pairs) to ornamented and unornamented caged female intruder. Pairs did not respond differently to female intruders showing some or no ornamentation. Bars denote means with plus/minus standard error lines overlaid.

We did not find any evidence for altered behavior of conspecifics or for a social cost in the context of territorial intrusions to females who acquired shoulder patches. Previous studies in females have found mixed support for social costs to female ornament expression. In Lovely Fairywrens, artificial enhancement of female plumage led to increased aggression of the signal bearer toward its own mirror image, indicating a social cost to dishonest signaling (Leitão et al. 2019). Conversely, in Pied Flycatchers, female decoys with a plumage signal on their head (white forehead patch) received fewer attacks than decoys lacking the signal, indicating no social costs to expression, and suggesting that the plumage patch signals fighting ability as it does in males (Morales et al. 2014). Our finding that females who acquired shoulder patches did not elicit a different territorial response from conspecific pairs is also interesting considering that altered conspecific response to manipulated signals is the proposed mechanism underlying observed transitions in behavior of the signal-bearer (reviewed in Vitousek et al. 2014; Webster et al. 2018). It is important to note that our trials with caged females did not mimic a natural situation, in that females of this species have not been observed invading neighboring territories on their own. Additionally, caged females may have been perceived as on a foray, as females of other fairywren species move into other territories to seek extra-territorial copulations by (Double and Cockburn 2000); we did not, however, observe any sexual displays by territory-holding males in response to these females, and the pair responding together is consistent with territory defense (eg. duetting and close approaches) rather than sexual functions of the response. We did not detect an effect of caged female overall activity level on the response of the territorial pair, though we may have failed to quantify subtle, but meaningful behavior of the caged female that affected the response.

Lastly, acquisition of shoulder patches by females did not affect the territorial response of mates. Male responses to STIs did not differ according either to their mate’s initial treatment (testosterone vs. control implant) or resulting plumage phenotype (ornamented vs. unornamented, Fig. S5). Therefore, we do not find evidence for the greater vocal territoriality in partially ornamented females being the product of greater aggression by their mates, or via compensating for lower aggression by their mates.

## Conclusions

We find support for testosterone facilitating the acquisition of a major component of a female ornament in a species with discrete variation in female ornamentation. Acquisition of ornamentation appears to be followed by enhanced vocal territoriality, consistent with the ornament functioning in the context of resource defense. In contrast to expectations, testosterone did not enhance vocal territoriality independently of ornamentation, as greater responses to STI occurred only after ornamentation was developed and not before. Our results are inconsistent with testosterone being an overall integrator of the ornamented female phenotype in White-shouldered Fairywrens. Rather, elevated testosterone appears to initiate a sequence of processes starting with the production of ornamental plumage signals, that leads to enhanced territorial behavior, but seemingly independent of testosterone. Our findings contribute to a growing body of research on the function of competitive female traits (Cain and Ketterson 2012; Tobias et al. 2012; Karubian 2013; Garamszegi 2014; Goymann and Wingfield 2014; Moreno et al. 2014; Cantarero and Cantarero 2015; Enbody et al. 2018; Leitão et al. 2019), and highlight the need for further experimental work testing the extent to which sexes share proximate mechanisms of phenotype expression.

## Acknowledgements

This study was supported by NSF grant (IOS-1354133 to J.K. and IOS-1352885 to H.S.), the Disney Worldwide Conservation Fund (to J.K.), the Washington State University Elling Fund and American Ornithological Society Research Grant (to J.B. and J.A.J.), the NSF Doctoral Dissertation Improvement Grant IOS-1701781 (to E.D.E.), and the Tulane University Ecology & Evolutionary Biology Department Student Research Grant and the Gunning Fund (to E.D.E and J.A.J.). We thank the Baivapupu Clan for granting us access to their land for this research. We’d like to thank I.R. Hoppe, M. Olesai, and K. Saiga and for assistance in the field. The National Research Institute in Papua New Guinea assisted with obtaining research permits for this work. Our experimental protocol was approved by IACUC (ASAF#04573) in the U.S. and the Center for Environmental Protection Agency in Papua New Guinea. M.S. Webster, A. Peters, and A. McQueen provided invaluable assistance with testosterone implant preparation. Finally, we thank P. Carter, E. Crespi, H. Watts, and the Center for Interdisciplinary Statistical Education and Research at Washington State University for helpful comments on experimental design and statistical analysis.

## Supplemental materials

**Table S1:** Summary of morphological results across study years; females separated by initial treatment and plumage type. Molt score and CP volume are means ± standard error.

| Treatment |  |  | Molt score |  |  | CP volume (mm<sup>3</sup>) |  |  |
| --- | --- | --- | --- | --- | --- | --- | --- | --- |
| Implant | Plumage | Individuals | Initial Molt | 7-11 Day Molt | 28+ Day Molt | Initial CP | 7-11 Day CP | 28+ Day CP |
| Control | Unorn. | 11 | 4.36 ± 0.94 | 4.50 ± 1.12 | 1.67 ± 0.92 | N | N | N |
| Testosterone | Unorn. | 5 | 2.40 ± 0.81 | 4.67 ± 1.86 | 4.17 ± 1.17 | N | 13.81 ± 8.62 | 3.76 ± 2.42 |
| Testosterone | Orn. | 15 | 5.27 ± 0.80 | 6.00 ± 0.88 | 6.13 ± 1.77 | N | 13.36 ± 4.00 | 0.78 ± 0.78 |

**Figure S1:**
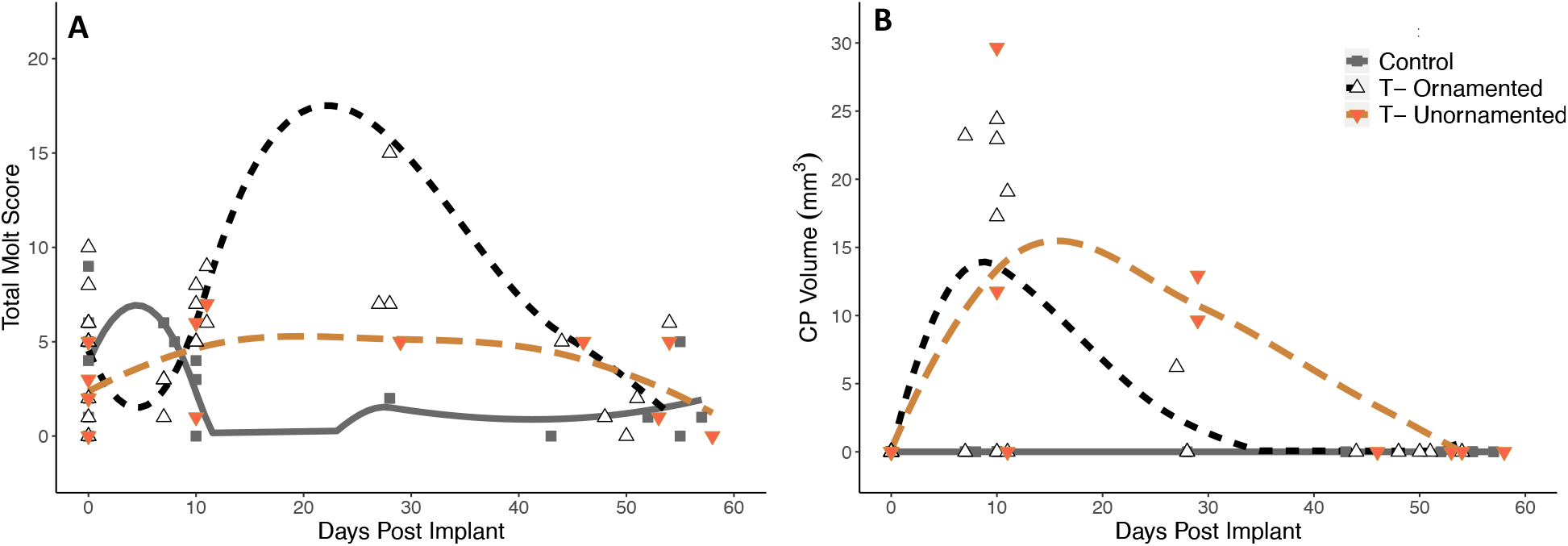
Total body molt scores (A) and cloacal protuberance volume (B) across capture days starting when implants were set (day 0) separated by initial treatment (T = testosterone) and plumage ornamentation (Ornamented = females with shoulder patches; Unornamented = females without shoulder patches). Each point is an individual capture for a female belonging to any of the 3 combinations of initial treatment and final plumage phenotype. Note absence of cloacal protuberances at capture beyond 40 days post-implant. One female was included from the testosterone-ornamented group who was sampled on day 27. Five females were sampled twice during the 28+ day period, each capture 27-23 days apart: one control, and two each from the testosterone-unornamented and testosterone-ornamented groups.

**Figure S2:**
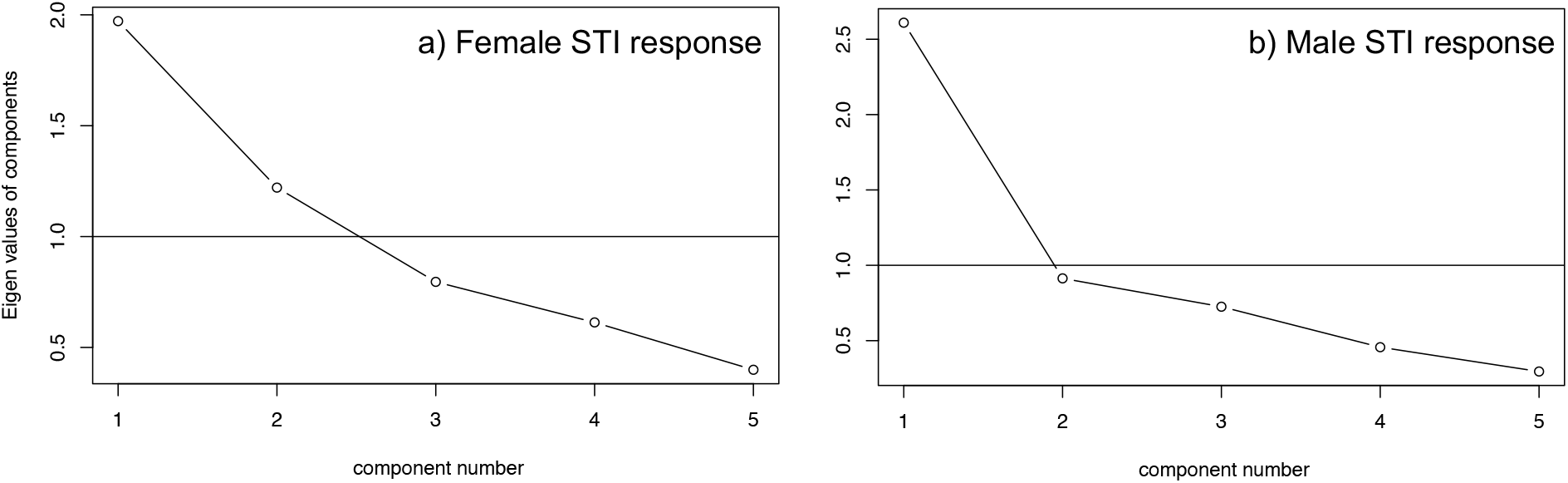
Scree plot for a) female and b) male STI response. Only components 1 and 2 were analyzed for females and component 1 for males.

**Figure S3:**
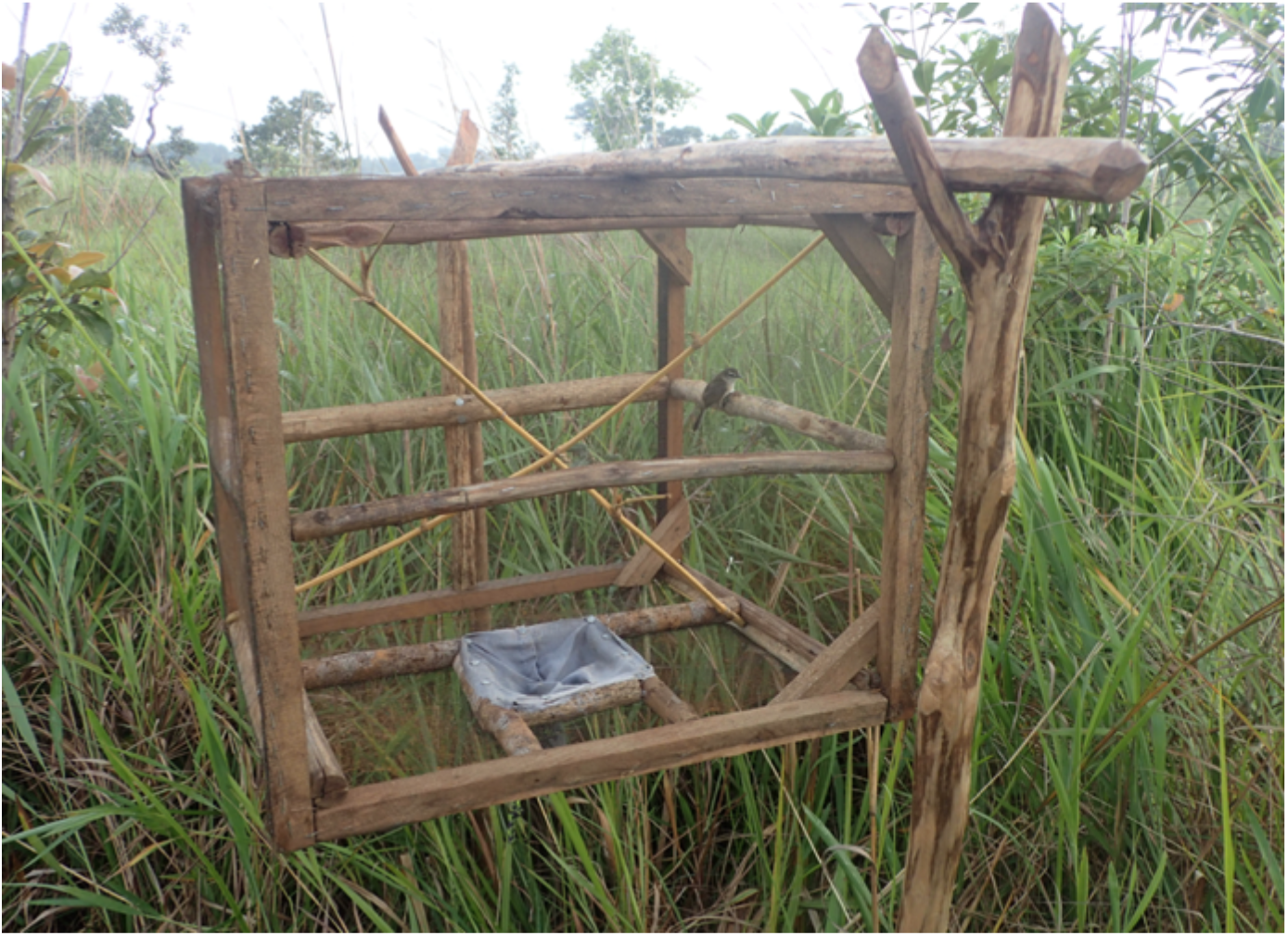
Cage trial setup with unornamented female inside.

**Figure S4:**
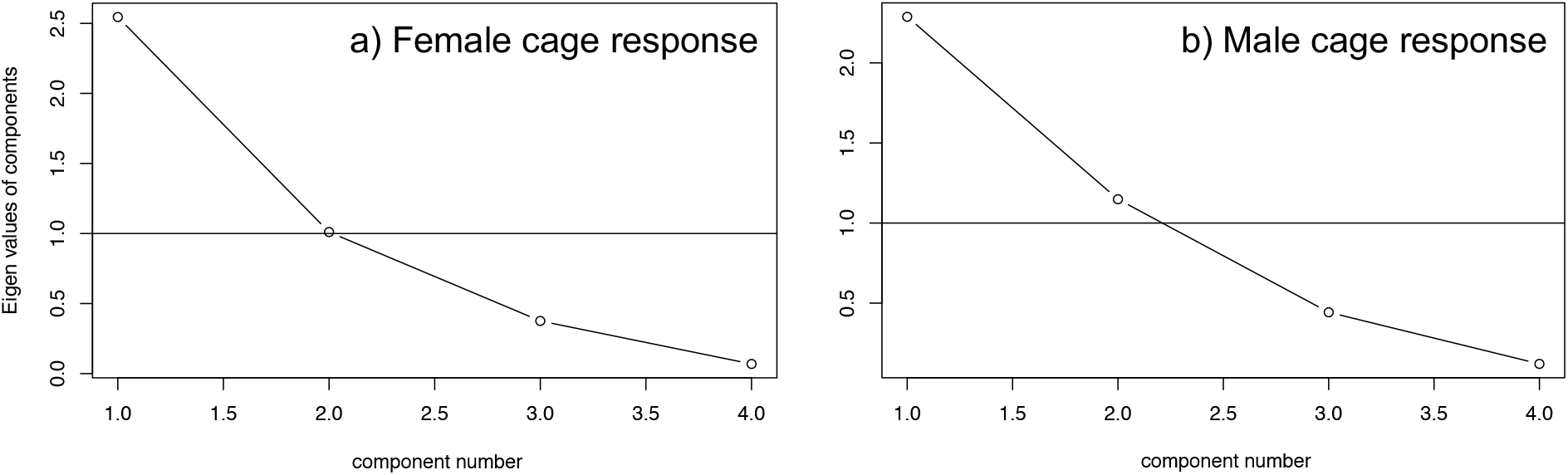
Scree plot for a) female and b) male response to ornamented or unornamented female presented in cages to territory holders. Only component 1 was further analyzed.

**Table S2:** Variable loadings for male response to simulated territorial intrusions

| Male STI Response | PC 1 |
| --- | --- |
| Standard deviation | 2.40 |
| Proportion of Variance | 0.48 |
| Latency to sing or duet | -0.70 |
| Latency to 5m | -0.74 |
| Song & duet rate | 0.77 |
| Time within 5m | 0.88 |
| Flyby rate | -0.05 |

**Figure S5:**
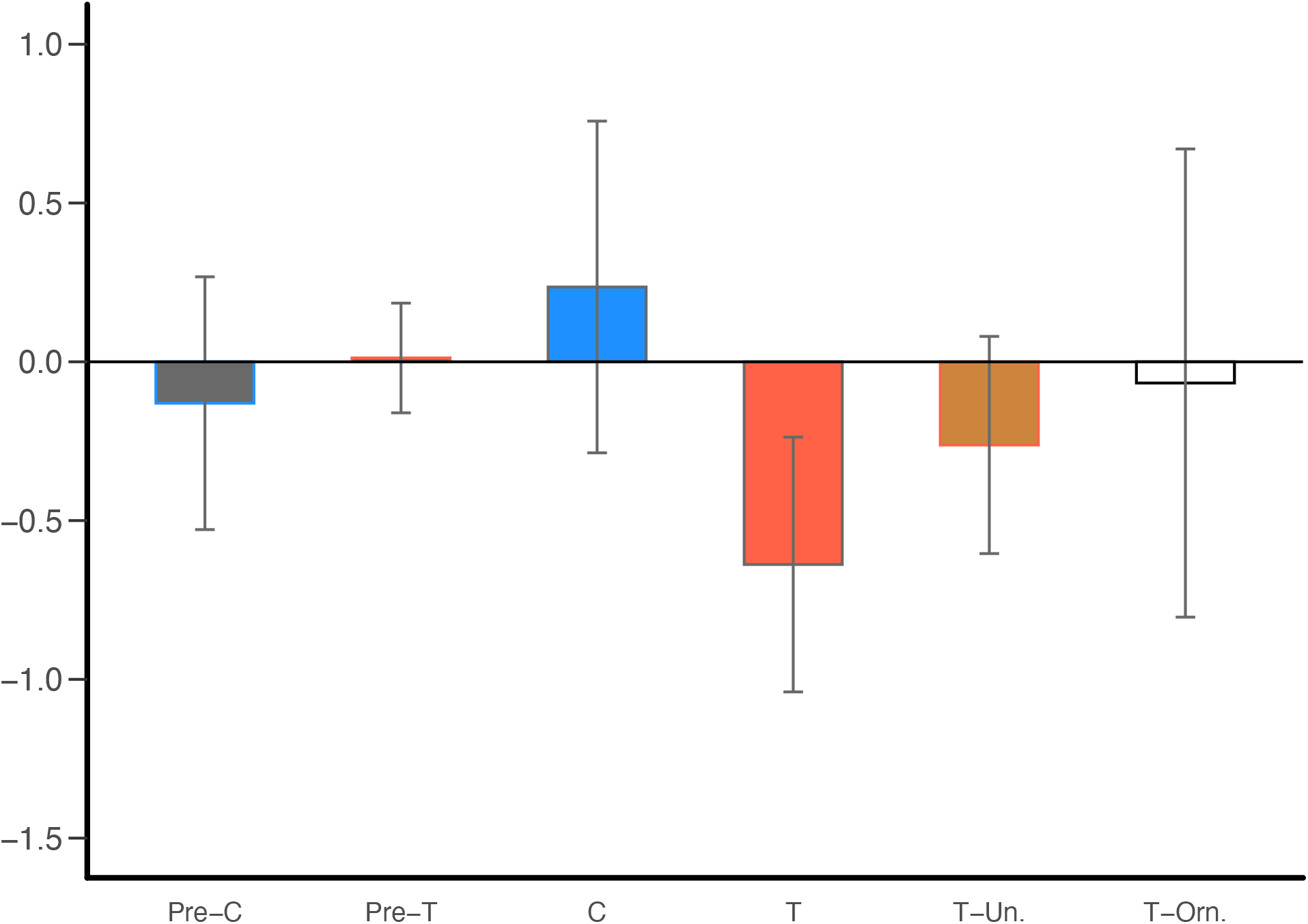
Male response to simulated territorial intrusions. We did not find a significant effect of the mate’s initial treatment or plumage phenotype on proximity to decoys or vocal response. Bars denote means with plus/minus standard error lines overlaid.

